# Use of CRISPR/Cas9 Gene Targeting to Conditionally Delete Glucocorticoid Receptors in Rat Brain

**DOI:** 10.1101/560946

**Authors:** Jessie R. Scheimann, Rachel D. Moloney, Parinaz Mahbod, Rachel L. Morano, Maureen Fitzgerald, Olivia Hoskins, Benjamin A. Packard, Yueh-Chiang Hu, James P. Herman

## Abstract

Glucocorticoid receptors (GR) have diverse functions relevant to maintenance of homeostasis and adaptation to environmental challenges. Understanding the importance of tissue-specific GR function in physiology and behavior has been hampered by near-ubiquitous localization in brain and body. Here we use CRISPR/Cas9 gene editing to create a conditional *GR* knockout in Sprague Dawley rats. To test the impact of cell-and region-specific *GR* deletion on physiology and behavior, we targeted *GR* knockout to output neurons of the prelimbic cortex.

Prelimbic deletion of *GR* in females caused deficits in acquisition and extinction of fear memory during auditory fear conditioning, whereas males exhibit enhanced active-coping behavior during forced swim. Our data support the utility of this conditional knockout rat to afford high-precision deletion of *GR* across a variety of contexts, ranging from neuronal depletion to circuit-wide manipulations, leveraging the behavioral tractability and enhanced brain size of the rat as a model organism.

## Introduction

The glucocorticoid receptor (GR) is a ligand (glucocorticoid)-activated transcription factor that participates widely in functions related to homeostasis and adaptation (1). Functional properties of GR action have been widely queried, using pharmacological, viral vector and electrophysiological approaches, with most of the literature using rat models (2-6). Recent decades have seen an increase in the use of mouse models to provide Cre-driver mediated deletion in specified cell populations (e.g. CaMKIIα neurons of the forebrain, Simpleminde-1 (Sim-1) neurons of the hypothalamus (7, 8)). While yielding valuable information on cell type-specific GR actions (9), the use of mice has proven problematic with regard to high-resolution behavioral and physiological analyses due to the different behavioral repertoire of mice (high strain and inter-experiment variability in memory tests) and the small size (e.g., prohibits repeated blood sampling)(10-13).

The emergence of CRISPR (Clustered Regularly Interspaced Short Palindromic Repeats)/ Cas 9-mediated gene editing now affords gene targeting in numerous species, including rat (14-16). For example, CRISPR/Cas9 methods allow for introduction of exon-flanking loxP sites to generate conditional knockout alleles and subsequent gene deletion following exposure to Cre recombinase, using either driver lines or viral vector approaches. In this study, we used CRISPR/cas9 to specifically insert loxP sites to sequences flanking exon 3 of the *Nr3c1* (*GR*) gene in Sprague Dawley rats (an outbred strain commonly used to explore GR function) via homology-directed repair (HDR). Our data provide validation of the *Nr3c1* gene editing in rats, and use viral vector-mediated Cre delivery to demonstrate targeting of GR deletion to specific cell types, brain regions and circuits. Functional efficacy of GR deletion in the prelimbic division (PL) of the medial prefrontal cortex (mPFC) (PL-PFC) was verified by sex-specific deficits in extinction of conditioned fear in females and a shift to active coping in the forced swim test (FST) in males. Overall, our study shows the utility of this gene editing technique to generate conditional gene deletion models that can leverage the considerable advantage of rats in behavioral and physiological research.

## Results

We directly manipulated the genome of Sprague Dawley rat zygotes by CRISPR/Cas9 to generate the floxed *Nr3c1* allele. We used a dual sgRNA strategy to delete the sequence containing exon 3 of the *Nr3c1* gene and repaired it with a donor plasmid that contains the deleted sequence, two flanking loxP sites, a right homologous arm at 2.57kb, and a left homologous arm at 1.95kb (Figure 1A, 1B). Because truncated sgRNAs increase targeting specificity (17), we chose six sgRNAs with various lengths (17-20 nt) (Supplementary Table 1) and validated their editing activity in rat C6 glioma cells by T7E1 assay (Supplementary Figure 1). We picked two sgRNAs, sg-2 and sg-6, for targeting the 5’ and 3’ sequences of exon 3, respectively. Both sgRNAs were 17nt in length, which is expected to provide high specificity (17). The two selected sgRNAs, Cas9 mRNA, and the donor plasmid were microinjected into ∼60 rat zygotes, followed by embryo transfer into pseudopregnant female rats. Seventeen pups were born. We identified that one of them (#60) was correctly targeted, which was confirmed by PCR with the external primers (Figure 1C, 1D) paired with the primers partially containing loxP sequences (P5-P6 and P10-P8 for 5’ and 3’ ends, respectively; Figure 1E, 1F). It was further confirmed by primer pairs P5-P7 and P9-P8, followed by ClaI and BamHI enzyme digestion and Sanger sequencing (data not shown). The fact that the offspring of rat #60 can be bred to homozygosity indicates the correct targeting of loxP sequences to the *Nr3c1* gene, instead of random integration (Figure 1G).

**Figure 1.**
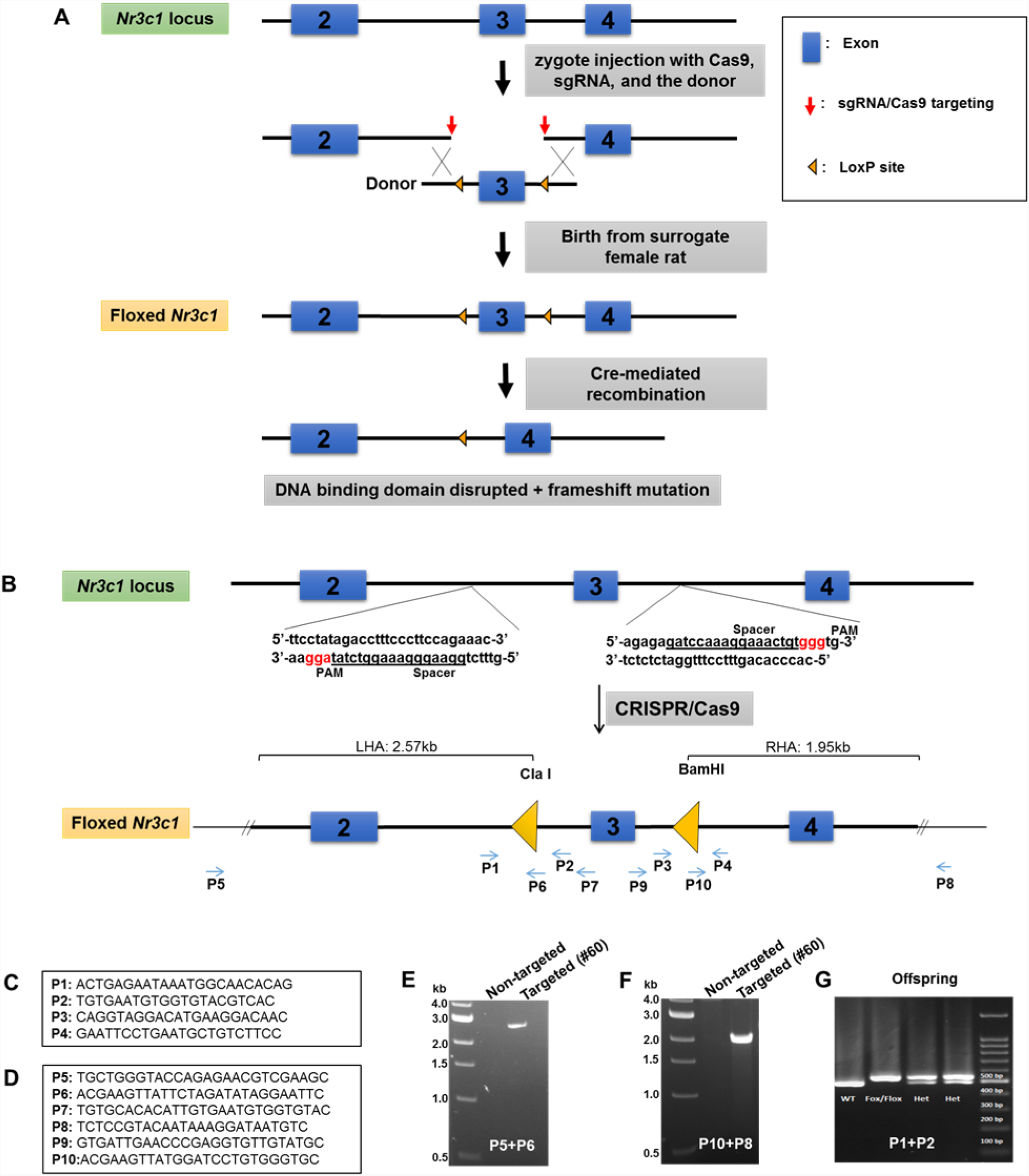
Generation of *Nr3c1* conditional knockout rat. (A) Schematic illustration of the targeting strategy. Two loxP sequences flanking exon 3 are inserted via CRISPR/Cas9-mediated deletion by two sgRNAs, followed by homology-directed repair with a donor plasmid. The CRISPR reagents were injected into the cytoplasm of rat zygotes, followed by embryo transfer into surrogate mothers for fetal development and birth. Correctly targeted rats were bred to homozygosity. Deletion of exon 3 is achieved by recombination of two loxP sequences upon the exposure of Cre recombinase. (B) The donor plasmid contains two loxP sequences flanking exon 3, restriction enzyme sites, and two homologous arms. The loxP sequence is located at the sgRNA cut site (3 nucleotides prior to the PAM) to block re-cutting. (C) Primers used to confirm the correctly targeted events are listed in the boxes. (D) Primers P5 and P8 are external to homologous arms. (E-G) Sample PCR results are shown.

### Validation of Conditional Gene Deletion: Viral Vector Targeting

To test the efficacy of this novel rat line, we administered adenoviral Cre recombinase constructs to drive regional, cell type-specific and projection-specific deletion of *Nr3c1*. Regional deletion targeted the basolateral amygdala (BLA), using human synapsin promoter-driven Cre recombinase (AAV8-hSyn-Cre, UNC Vector Core, NC, USA) microinjections. Cre-positive cells were devoid of nuclear GR immunoreactivity in SD:nr3c1^fl/fl^ rats (Figure 2A), whereas the vast majority of Cre-positive cells co-expressed GR in wildtype controls (SD:nr3c1^wt^) (Figure 2B) injected with the same viral construct (AAV8-hSyn-Cre, UNC Vector Core).

**Figure 2.**
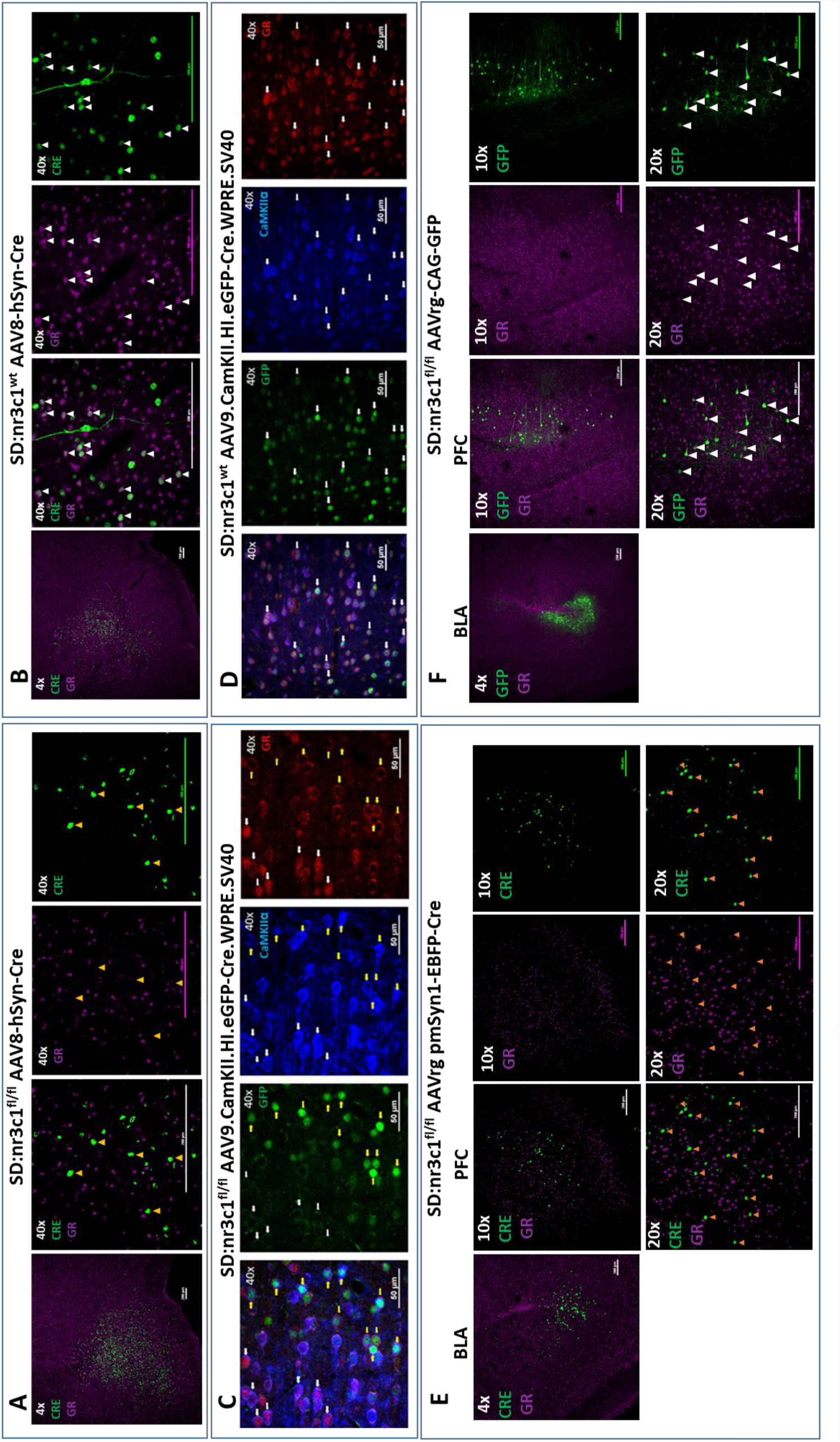
Viral Validation of *Nr3c1* conditional knockout rat. (A) AAV8-hSyn-Cre administration to the basolateral amygdala (BLA) in SD:nr3c1^fl/fl^ rats (panel A1-site of injection, 4X). Note the absence of GR (purple) in Cre^+^ neurons (green) (panels A2-4, 40x, yellow triangles). (B) AAV8-hSyn-Cre administration to the basolateral amygdala (BLA) in SD:nr3c1^wt^ rats (panel B1-site of injection, 4X). Note the presence of GR (purple) in Cre^+^ neurons (green) (panels B2-4, 40x, white triangles). (C) AAV9.CamKII.HI.eGFP-Cre.WPRE.SV40 (CaMKIIα Cre) injected into SD:nr3c1^f/f^ rats results in decreased GR (C4 yellow arrows) in cells infected with virus as shown by GFP labeling (C2 yellow arrows) that are also CaMKIIα positive (C3 yellow arrows). CaMKIIα cells not infected with GFP, show endogenous GR expression (C1-4 white arrows). (D) AAV9.CamKII.HI.eGFP-Cre.WPRE.SV40 (CaMKIIα Cre) injected into SD:nr3c1^wt^ rats (panel C2-4 40x white arrows) shows endogenous GR staining (D4) in cells infected with virus shown in GFP (D2) and CaMKIIα positive (D3) (E) AAVrg-pmSyn1-EBFP-Cre administration to the BLA in SD:nr3c1^fl/fl^ rats (panel E-1 site of injection, 4X) and retrograde trafficking of the virus to cell somas in the prefrontal cortex (PFC) (panel E2-4). Note absence of GR expression (purple) in Cre^+^ neurons (green) (panel E5-7, 20x, yellow triangles). (F) AAVrg-CAG-GFP administration to the BLA in SD:nr3c1^fl/fl^ rats (panel F1-site of injection, 4X) and retrograde trafficking of the virus to cell somas in the PFC (panel F2-4, 10X). Note expression of GR (purple) in GFP^+^ neurons (green) (panel F5-7, 20x, white triangles).

We then assessed cell-type specific deletion by injection of AAV9.CamKII.HI.eGFP-Cre.WPRE.SV40 (AAV-CaMKIIa-Cre) virus into the prefrontal cortex of SD:nr3c1^wt^ or SD:nr3c1^fl/fl^ rats. Infusions of AAV-CaMKIIa-Cre caused widespread loss of GR immunoreactivity within GFP positive cells in the PFC of AAV-CaMKIIa-Cre virus injected SD:nr3c1^fl/fl^ rats (figure 2C yellow arrows), but no GR loss is noted in wildtype control (SD:nr3c1^wt^) injected rats (figure 2D white arrows) or in uninfected CaMKIIα cells in SD:nr3c1^fl/fl^ rats (figure 2C white arrows). Specificity of deletion of GR was confirmed in CaMKIIα positive neurons in AAV-CaMKIIa-Cre injected animals, with the vast majority of CaMKIIα immuno-positive cells showing deletion of GR (figure 2D).

Finally, we used an intersectional approach to examine connectional deletion of GR, focusing on PFC projection neurons to the BLA. Retrogradely-infected (Cre-positive) neurons were observed in the PFC after administration of AAVrg pmSyn1-EBFP-Cre, (Addgene) to the BLA. We did not observe GR immunoreactivity in SD:nr3c1^fl/fl^ rats (Figure 2E), whereas substantial proportions of PFC-BLA projecting neurons contained GR immunoreactivity in control virus (AAVrg-CAG-GFP, Addgene) injected animals (Figure 2F). These studies highlight the novel use of this rat model to query not only the role of GR in a specific region in isolation but also how GR functions as part of an integrated circuit.

### Behavioral Consequences of Targeted GR Deletion; Implications for Fear and Coping Behaviors

We next performed a functional test of viral Cre-mediated GR knockout, focusing on the PL-PFC. SD:nr3c1^fl/fl^ (GRKO) rats and wild type littermate controls SD:nr3c1^wt^ (Control) all received injection of AAV9.CamKII.HI.eGFP-Cre.WPRE.SV40 (CaMKIIα Cre) [Penn Vector Core] into the PL-PFC (Figure 3C). The PFC plays a critical role in extinction of emotional memory (e.g. conditioned fear), selection of emotional coping strategy and HPA axis reactivity (18-21). CaMKIIα is a calcium binding protein that is most commonly found in glutamatergic neurons of the forebrain, thus we are targeting the excitatory output of the PL-PFC. We therefore tested whether GR knockout in CaMKIIα targeted neurons in this region affected extinction of conditioned fear to an auditory fear conditioning paradigm, and behavioral coping during the FST. For fear conditioning, rats were exposed to 5 tone shock pairings on the first day (acquisition), followed by 2 days of 20 tones without a paired shock, (extinction and extinction recall). Data was binned for clarity into 5 tones per bin. Freezing during the tones was measured as fear behavior. Freezing in female rats with PL-PFC targeted GRKO (SD:nr3c1^fl/fl^ plus CaMKIIα Cre) increased in acquisition [GRKO F(1,52) = 5.553; p = 0.035; wt mean = 40.784 SEM = 4.038 n = 10; GRKO mean = 57.264 SEM = 5.710 n = 5; pη^2^ = 0.043], but there was no significant interaction effect[GRKO x tone F(4,52) = 1.292; p = 0.285; pη^2^ = 0.024] (figure 3A). Extinction of fear conditioning was delayed in the female GRKO group relative to controls, as indicated by a significant time x GRKO interaction effect [GRKO x time F(3,39) = 4.184; p = 0.012; wt mean = 48.174 SEM = 6.497 n = 10; GRKO mean = 64.157 SEM= 9.188 n = 5; pη^2^ = 0.043] (figure 3A). Extinction recall was also impaired in female PL-PFC GRKO rats relative to controls (significant time x GRKO interaction) [GRKO x time F(3,31) = 3.796; p = 0.020; wt mean = 32.484 SEM = 7.721 n = 10; GRKO mean = 55.166 SEM = 8.458 n = 5; pη^2^ = 0.046] (figure 3A). Meanwhile, males with PL PFC targeted GRKO (SD:nr3c1^fl/fl^ plus CaMKIIα Cre) did not differ from controls [Acquisition GRKO x tone F(1,79] = 1.827; p = 0.136; wt mean = 6.739 SEM = 0.49 n = 10; GRKO mean = 6.080 SEM = 0.580 n = 6; pη^2^ =0.036] [Extinction GRKO x tone F(1, 63) = 0.873; p = 0.463 wt mean = 64.112 SEM = 6.701 n = 10; GRKO mean = 57.107 SEM = 8.651 n = 6; pη^2^ = 0.01] [Extinction Recall GRKO x tone F(1,63) = 0.592; p = 0.624; wt mean = 56.236 SEM = 5.163 n = 10; GRKO mean = 51.532 SEM 6.665 n = 6; pη^2^ = 0.014] (figure 3B). While most lesion and inactivation studies have shown that the PL PFC is critical for appropriate fear responding (19-21), the use of our new GR floxed rat model has shown that GR in the glutamatergic excitatory output of the PL-PFC may be more critical for female expression and extinction of conditioned fear, and less so for males.

**Figure 3.**
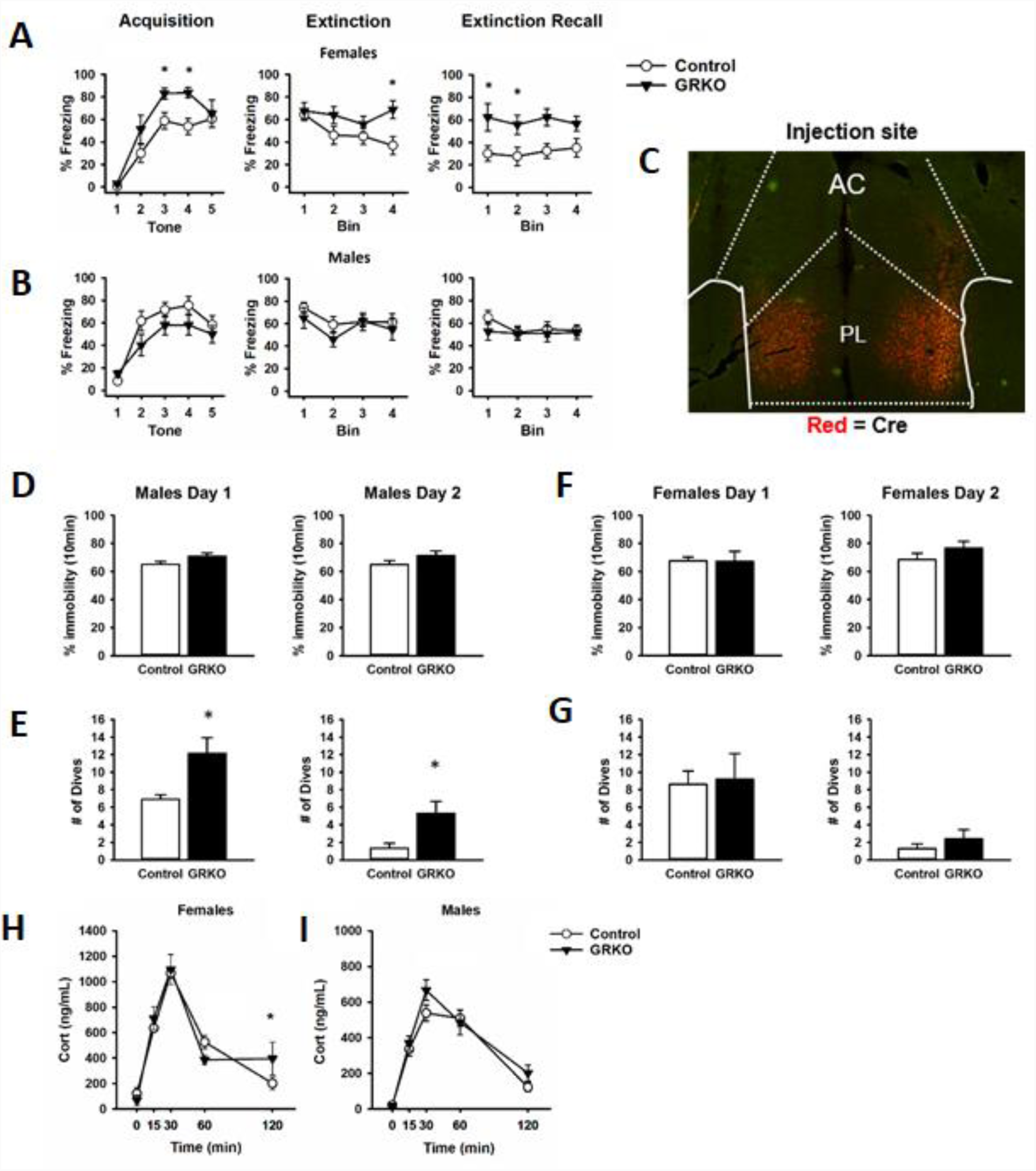
Behavioral profile with *Nr3c1* knockout in CaMKIIα cells in the Prefrontal Cortex. A) Female rats with PFC-PL CaMKIIα GRKO showed heightened freezing to tone shock pairings compared to controls, as well as deficits in fear extinction, and heightened freezing during extinction retrieval indicating an inability to extinguish conditioned fear. B) Male rats with PFC-PL CaMKIIα GRKO did not differ from controls during auditory fear conditioning. C) Representative image of CaMKIIα injection site and area of infection in the Prelimbic cortex of the PFC. Red = Cre recombinase protein. Male rats with CaMKIIα GRKO in the PL PFC did not show differences from controls in immobility on either day in the forced swim test D), but CaMKIIα male rats did show a greater number of dives on both days E) suggesting a more active coping behavior. F-G) Female rats with CaMKIIα GRKO did not show differences from controls in the forced swim test. H) Females with PFC-PL CaMKIIα GRKO show higher corticosterone release to acute restraint at the 120min time point as well as a significant GRKO x time interaction. I) While male rats with GR knockout in the PL PFC CaMKIIα GRKO do not show differences from control in corticosterone release to acute restraint. * = p<0.05. (GRKO = SD:nr3c1^fl/fl^ plus AAV CaMKIIα Cre), n = 5-6 male and female; Control = SD:nr3c1^wt^ plus AAV CaMKIIα Cre, n = 10 male and female).

The PFC has also been implicated in coping behaviors and behavioral adaptation during acute and chronic stress. We used the FST to investigate remodeling of behavioral coping strategy during an acute stress challenge. There was no effect of PL-PFC-driven GRKO (SD:nr3c1^fl/fl^ plus CaMKIIα Cre) on immobility time in the FST on either day 1 or day 2 for male rats [Day 1 immobility T(14) = −1.8.48; p = 0.085; wt mean = 65.167; SEM = 2.051; GRKO mean = 71.003 SEM = 2.199 n = 6; 95% confidence interval = −12.608 to 0.937; Cohen’s d = 0.983] [Day 2 immobility T(14) = −1.411; p = 0.180; wt mean = 65.052 SEM = 2.819 n = 10, GRKO mean = 71.341 SEM = 3.306 n =6; GRKO mean = 428.049 SEM = 21.726 n = 6; 95% confidence interval = −15.849 to 3.269; Cohen’s d = 0.739] (figure 3D). However, males with CaMKIIα-driven PL-PFC GRKO (SD:nr3c1^fl/fl^ plus viral CaMKIIα Cre) increase diving frequency on day 1 [T(14) = 3.252; p = 0.005; wt mean = 6.900 SEM = 0.547 n = 10; GRKO mean = 12.167 SEM = 1.922 n = 6; 95% confidence interval = 1.793 to 8.741; Cohen’s d = 1.64] (figure 3E) relative to controls, consistent with an altered coping strategy. This behavior persisted into the second day of the FST test, with significantly higher number of dives for males with PL-PFC GRKO (SD:nr3c1^fl/fl^ plus viral CaMKIIα) [T(14) = −3.468; p = 0.003; wt mean = 1.100 SEM = 0.407 n = 10; GRKO mean = 5.333 SEM = 1.453 n =6; 95% confidence interval = −6.852 to −1.615; Cohen’s d = 1.747] (figure 3E). Females with PL-PFC GRKO (SD:nr3c1^fl/fl^ plus CaMKIIα Cre) did not differ from controls in total immobility (figure 3F) or number of dives (figure 3G) [Day 1 Immobility = T(13) = 0.077; p = 0.939; wt mean = 67.691 SEM = 2.468 n = 10; GRKO mean = 67.221 SEM = 7.168 n =5; 95% confidence interval = −12.558 to 13.497; Cohen’s d = 0.039] [Day 2 immobility = T(13) = −1.083; p = 0.298; wt mean = 68.491 SEM = 4.621 n = 10; GRKO mean = 76.691 SEM = 5.302 n = 5; 95% confidence interval = −24.557 to 8.158; Cohen’s d = 0.619] [Day 1 dives = T(13) = −0.187; p = 0.854; wt mean = 8.6, SEM = 1.607; GRKO mean = 7.294, SEM = 3.262; 95% confidence interval = −7.525 to 6.325; Cohen’s d = 0.096] [Day 2 dives = T(13) = −1.316; p = 0.211; wt mean = 2.346 SEM = 1.077 n = 10; GRKO mean = 6.459 SEM = 3.991 n = 5; 95% confidence interval = −10.864 to 2.637; Cohen’s d =0.304 (data transformed to square root to pass equal variance]. Although there were no changes in total immobility in the FST, increased diving in the males with PFC-PL GRKO suggests GR in excitatory neurons of the PL-PFC may facilitate a shift, selectively in males, to more diverse active escape behaviors than just swimming and climbing alone.

Our lab has previously shown that GRKO in the PL-PFC increases corticosterone to acute restraint (2). We subjected male and female GRKO and control rats to an acute restraint challenge and took tail blood samples at 0min, 15min, 30min, 60min, and 120min from the start of restraint to measure plasma corticosterone. There was a GRKO x time interaction [F(4,66) = 3.020; p = 0.027; wt mean = 510.345 SEM = 34.599 n = 10; GRKO mean = 518.022 SEM = 55.460 n = 5; pη^2^ = 0.024] in females, with post hoc analysis revealing delayed shut-off (120 min time point) of the HPA axis stress response in PFC-GRKO rats [p = 0.050] (figure 3H). There was no effect of GRKO or GRKO/time interaction observed in males [F(1,64) = 2.046; p = 0.180; wt mean = 305.569 SEM = 18.445 n = 10; GRKO mean = 348.116 SEM = 23.332 n = 6; pη^2^ = 0.012] (figure 3I). The data suggest that female SD:nr3c1^fl/fl^ rats with PL-PFC GRKO may have negative feedback inhibition due to the inability to return to a similar resting level of corticosterone as control rats. The lack of differences after PL-PFC GRKO in the males compared to our previous study may be due to the specificity of our virus to knockdown only in CaMKIIα cells, whereas previously a ubiquitous promotor on a lentiviral delivered shRNA was used which targets all neuronal cell types, including inhibitory GABAergic interneurons (2).

## Discussion

Prior studies document generation of knockout and Cre dependent knockout rat models using CRISPR/Cas9 as a molecular tool (15, 22) reviewed in (14). Here we created a conditional (Cre-recombinase dependent) GR knockout rat using CRISPR/Cas9. Validation of genome integration was accomplished by PCR, and Cre-dependent deletion by site and circuit specific viral vector expression of Cre. Finally, circuit specific deletion was used to document GR-dependent modification of behavioral and neuroendocrine function by the prelimbic cortex. The CaMKIIα promoter is thought to largely direct Cre expression to cortical projection neurons with over 80% efficacy (23) and thus it is likely our manipulations were able to successfully target excitatory projection neurons.

Deficits in fear conditioning and alterations in behavioral coping style are a common phenotype in murine GR knockout models as well as following chronic stress (2). We used site and cell specific knockout of GR to demonstrate that CaMKIIα cells in the PL-PFC require GR for appropriate fear responses and extinction in females. Conversely, male rats lacking GR in the PL-PFC, shift to an active escape behavior in the FST, suggesting adoption of an active coping strategy. Females (but not males) evidenced deficient shut-off of HPA axis stress responses, consistent with a sex-specific dependence on the PL GR for full feedback inhibition. The data highlight a strong interaction between GR signaling and sex in coordination of prefrontal cortical signaling mechanisms. Moreover, the data provide functional evidence that CRISPR/Cas9 can be used to provide high-resolution cell and site specific assessment of GR action in brain, leveraging the advantages of the rat as a model organism.

Prior studies have used promoter-specific Cre driver mouse lines to generate targeted GR knockout in mice. Mouse studies demonstrate that CaMKIIα-Cre directed GR deletion in the forebrain (targeting exon 2 or 3) promote anxiety-related behavior, passive coping and corticosterone hypersecretion in male, but not female mice (9, 24, 25). It is important to note that the mouse CaMKIIα driver line deletes GR in multiple brain regions, including cortex, hippocampus, basolateral amygdala, caudate and bed nucleus of the stria terminalis, many of which interface with stress and emotional behavior. The extensive knockout makes it impossible to specify circuit-specific roles of stress hormone signaling in behavior and stress physiology. Prior studies have not specified GR-specific deficits in cognitive behaviors, which can be difficult to assess in mouse models (10-12). Here, we show that by using these viral constructs with specific promoters (CaMKIIα) in GR floxed rats, we can not only drive expression of Cre-recombinase in a cell-type specific manner, but also a defined region, allowing precise investigation of the role of GR in behavior.

There is a possibility of genetic modifications, even with the highly site specific editing of the CRISPR/Cas9 system, when inserting LoxP sites into the genome (reviewed in (26-28)). We tested the most likely frameshift insertion and deletion (indel) mutations that could be introduced by unexpected non-homologous end joining or inaccuracies in the sgRNA targeting. We used PCR amplification and sequencing to determine that there were no mismatches in these sequences compared to wildtype controls (Supplementary Table 2&3). We are thus confident that our LoxP insertion did not cause any genetic mutations that would interfere with *in vivo* studying of the GR functioning in the rats on a genetic basis. As an additional control, we further investigated physiological and behavioral effects of this targeted insertion. We performed multiple tests of behavior, and observed no effect of gene targeting on any behavioral endpoint (Supplementary Figure 2). Furthermore, we found no differences in bodyweight (Supplementary Figure 3) or organ weights commonly changed with chronic stress (Supplementary Table 4).

Our functional studies indicate that female (but not male) rats with knockout of GR in the PL-PFC have heightened fear response and impaired extinction. Importantly, these data inform prior PL-PFC lesion/inactivation studies, which indicate that the PL is critical for appropriate encoding and expression of conditioned fear (20, 29, 30). The data indicate that GR signaling is an important component of this process in females, but may be less so in males (although the increased freeze times in males may reflect a more ‘intense’ response to the shock, which may result in ceiling effects on subsequent exposure to cues). In the FST, males but not females increased the number of diving events. As diving is considered an active coping behavior (31-33), it is likely that GR in the PL-PFC also interfaces with coping behaviors in a sex-specific manner. In combination, these data indicate that use of enhanced precision methods of gene targeting such as CRISPR/Cas9 in rat reveal substantive new information on the biology of the PL-PFC and its interaction with biological sex.

Deficits in conditioned fear and glucocorticoid feedback efficacy are associated with depression, post-traumatic stress disorder and other stress related diseases in human. It is important to consider that these disease states are overrepresented in women (2;1), and also involve modification of prefrontal cortical circuitry in neuroimaging studies (34-37). Emergence of behavioral and neuroendocrine deficits following targeting of GR in the PL of females may reflect a mechanism underlying selective vulnerability of females to stressful life events, suggesting that appropriate GR signaling is required to mitigate the impact of adversity.

Overall, we demonstrate that CRISPR/Cas9 gene editing is effective in generating novel tools for bio-behavioral research in a highly tractable model organism with a long and well-documented history. As noted in our functional studies, the SD:nr3c1^fl/fl^ line can be of significant value in the context of higher order behavioral assays (cognitive behaviors, goal-directed behaviors, reward behaviors/drug self-administration) that can be often difficult to implement and/or interpret in other rodent organisms, such as mice. With the growing number of viral Cre constructs and the recent surge in development of rat Cre driver lines, the CRISPR/Cas9 method can provide site and cell type gene deletions or manipulations that are vital for our understanding of mechanisms of stress pathologies and stress-related diseases.

## Methods

### Generation of the floxed *Nr3c1* line

Six sgRNAs targeting the sequences flanking exon 3 of *Nr3c1* were designed using the CRISPR Design Tool website (http://www.genome-engineering.org/). The complementary oligos (IDT) with overhangs were cloned into the BbsI site of the pX459 vector (Addgene #48139), according to the published methods (16). Editing activity was validated by the T7E1 assay (NEB) in rat C6 glioma cells (ATCC), compared side-by-side with *ApoE* sgRNA that was previously shown to work efficiently in rat zygotes (22). Validated sgRNA were *in vitro* transcribed by MEGAshorscript T7 kit and then purified by MegaClear kit (ThermoFisher). *Cas9* mRNA was *in vitro* transcribed by mMESSAGE mMACHINE T7 ULTRA kit (ThermoFisher), according to manufacturer’s instructions. Two sgRNAs (50 ng/ul ech), *Cas9* mRNA (100 ng/ul), and the donor plasmid (100 ng/ul) were mixed in 0.1X TE buffer and injected into the cytoplasm of one-cell-stage embryos of Sprague Dawley genetic background rats via a piezo-driven cytoplasmic microinjection technique. Injected embryos were immediately transferred into the oviductal ampulla of pseudopregnant females. Live born pups were genotyped by PCR, enzyme digestion, and Sanger sequencing. Rats were bred and housed in a vivarium with a 12-hour light/dark cycle. All animal studies were approved by the Institutional Animal Care and Use Committees of the Cincinnati Children’s Hospital Medical Center and University of Cincinnati.

### Breeding and Genotyping

A female Sprague Dawley rat (#60) containing the floxed Nr3c1 alleles was generated by the Transgenic Animal and Genome Editing Core Facility (Figure 1A, 1B). The founder rat that was heterozygous for the loxP knock-in sequences was crossed with a WT Sprague-Dawley male. F1 heterozygous offspring were bred to generate F2 and F3 offspring. We used F2 and F3 heterozygote animals to generate F/F [SD:nr3c1^fl/fl^] or WT [SD:nr3c1^wt^] littermate controls for behavioral and molecular experiments. Our breeding scheme was designed to minimize inbreeding by avoiding sibling x sibling mating.

We used four different sets of primers (Figure 1C, 1D) to verify that the loxP sequences were inserted in the 5’ and 3’ sides of exon 3. DNA was extracted from tail blood sample by using PureLink^®^ Genomic DNA Kits cat N: K1820-01, K1820-02, K1821-04. To discriminate the genotype of the rats, we performed PCR reactions with the primers that are listed in Figure 1C, 1D, by using FailSafe™ PCR 2X PreMix D CatN: FSP995D, and Dream Taq enzyme Cat N: EP0701 in PTC-200 Peltier Thermal Cycler. All the PCR product was analyzed with electrophoresis agarose gel and the images were captured by Axygen^®^ Gel Documentation System (Figure 1E-G).

For confirmation by sequencing, the expected bands from homozygote rats and littermate WT control were separated from the agarose gel and purified with Thermo Scientific™ GeneJET™ Gel Extraction Kit. The purified DNA was sent to Cincinnati Children’s Hospital Medical Center (CCHMC) DNA Sequencing and Genotyping Core to sequence the 3’ and 5’ sites of exon 3 of the *Nr3c1* gene, using the P1-P2 and P3-P4 primers. The sequencing data clearly showed the inserted flox sequences in both 3’ and 5’ sides of *Nr3c1* in Flox/Flox rats. The flox sequences were not seen in the WT littermate controls (Figure 1G).

### Stereotaxic surgery

Adult male and female SD:nr3c1^fl/fl^ and SD:nr3c1^wt^ rats (350g) were singly housed on a 12hr light/dark cycle in a temperature-and humidity-controlled housing facility at the University of Cincinnati. All experimental procedures were conducted in accordance with the National Institutes of Health Guidelines for the Care and Use of Animals and were approved by the University of Cincinnati Institutional Animal Care and Use Committee. Animals were deeply anaesthetized with 4-5% Isoflurane, prior to placement in the stereotaxic frame (Kopf Instruments) and sedation maintained at 2-3% isoflurane during surgery. A 2ul Hamilton syringe was used to administer viral constructs. The needle was gently lowered to the predefined coordinates for BLA (AP: −2.7, ML:+/- 4.8, DV: 8.8) or PL PFC (AP:+3.0, ML: +/-0.6, DV:-3.3) and a 5 minute rest period was observed. The virus was infused over 5 minutes (1ul/5mins). After infusion the needle remained in place for a further 5 minutes. The needle was slowly removed and the hole sealed with gelfoam. After completion of all infusions, the surgical site was closed with surgical staples and animals were singly housed for recovery.

### Viral constructs

AAV8-hSyn-Cre (titer: 6.5×10^12^ molecules/ml) was sourced from the UNC Vector Core (Chapel Hill, NC, USA). AAVrg-CAG-GFP (titer: 5×10^12^ vg/mL, this construct was a gift from Edward Boyden to Addgene-viral prep # 37825-AAVrg) and AAVrg pmSyn1-EBFP-Cre (titer: 6×10^12^ vg/mL, this construct was a gift from Hongkui Zeng to Addgene-viral prep # 51507-AAVrg (38)), were sourced from Addgene (MA, USA). All constructs were administered 1ul bilaterally and a minimum of 3 weeks incubation was allowed.

For CaMKIIα cell-specific knockout and fear conditioning and forced swim studies, 0.1ul of AAV9.CamKII.HI.eGFP-Cre.WPRE.SV40 (CaMKIIα Cre) [Penn Vector Core] (titer: 6.544×10^13^ diluted to 6.544×10^11^] was injected bilaterally and allowed 5 weeks to incubate.

### Immunohistochemistry

3-4 weeks after viral injection animals were injected with an overdose of pentobarbital and perfused transcardially with 0.1M PBS until blood was clear followed by 4% paraformaldehyde/0.1M PBS for 15 min. The brains were postfixed overnight at 4C in 4% paraformaldehyde/0.1M PBS, rinsed 2×3 times with 0.1M PBS and cryoprotected in 0.1M PBS containing 30% sucrose +.01% sodium azide at 4C until the brains sank in the sucrose solution. Brains were frozen to −20C on the stage of a sliding microtome and sectioned at 35μm, collected in a series of 6 and placed in cryoprotectant consisting of 10% polyvinyl-pyrrolidone-Mol Wt 40,000, + 500 ml of 0.1M PBS + 300 ml of ethylene glycol + 30 % sucrose and.01% sodium azide. A full series (6) of sections was labeled for GFP, GR and Cre Recombinase immunohistochemistry. The sections were processed as follows: 1% sodium borohyhdride/0.1M PBS for 30 min, rinsed 6×5 min −0.1M PBS, 1% hydrogen peroxide/0.1M PBS −10 min, rinsed 6×5 min-0.1M PBS, rinsed additionally in 0.1M PBS 4×15 min. Sections were blocked by incubation 0.1M PBS containing 4% goat serum with 0.4% Triton X-100 and 0.2% BSA for 2 hours, followed by a cocktail including polyclonal rabbit GR antibody (Santa Cruz-Cat# sc 1004) diluted 1:500, monoclonal mouse Cre Recombinase antibody (Millipore Cat # MAB 3120) diluted 1:1000 and no GFP antibody label (native virus expression) in blocking solution consisting of 0.1M PBS containing 4% goat serum with 0.4% Triton X-100 and 0.2% BSA overnight at RT. After overnight antibody incubation, sections were rinsed 0.1M PBS 3×5 min, then incubated in a cocktail of goat-anti-mouse CY3 conjugated IgG (Invitrogen-Cat#A32727) diluted 1:800 and goat anti rabbit CY5 conjugated IgG (Invitrogen-Cat # A32733) diluted 1:800 in 0.1M PBS for 45 min, and rinsed 4×5 min in 0.1M PBS. All sections were mounted and viewed on a Nikon C2 Plus Confocal Microscope.

For AAV9.CamKII.HI.eGFP-Cre.WPRE.SV40 (CaMKIIα Cre) cell-specific knockout, brains were processed the same until immunohistochemistry. The sections were processed as followed: slices rinsed 5×5 in 0.1M PBS, incubation in 1.5% 10mM Trisodium Citrate in PBS at 80°C for 10min, rinsed 5×5 0.1M PBS, blocked in 0.2% BSA + 4% goat serum 1hr room temperature, and then incubated overnight at 4°C in a cocktail of block plus rabbit polyclonal GR antibody (Invitrogen PA1-511A) diluted 1:800, chicken polyclonal GFP antibody (abcam ab13970) diluted 1:2000, and mouse monoclonal CaMKII antibody (abcam ab22609) diluted 1:250. The following day sections were washed 5×5 in 0.1M PBS and incubated at room temperature for 1 hr in the following secondary antibodies diluted 1:500 in blocking solution: goat-anti-Rabbit Cy3 conjugated IgG (Invitrogen A10520), goat-anti-chicken conjugated Alexa 488 (Invitrogen or A11039), and goat-anti-mouse CY5 conjugated IgG (Invitrogen A10524).

Verification of virus placement for functional PL-PFC knockdown study (AAV9.CamKII.HI.eGFP-Cre.WPRE.SV40) was completed via immunohistochemical analysis of Cre recombinase protein. Sections (series of 6) were processed as stated above before immunohistochemistry. Sections were washed 5×5 in 0.1M PBS and blocked in 0.2%BSA + 0.4% Triton-X 100 + 4% goat serum. Sections were then incubated overnight in block at 4°C in monoclonal mouse cre recombinase antibody (Millipore MAB 3120) diluted 1:1000. The following day sections were washed 5×5 in 0.1M PBS and incubated in goat-anti-mouse CY3 conjugated IgG (Invitrogen A32727) diluted 1:500 at room temperature for 1hr. All sections were mounted and viewed on a Nikon C2 Plus Confocal Microscope.

### Corticosterone measurements after acute restraint

Rats were restrained in a well ventilated plastic restrainer for 30min. Blood samples were taken via tail nick (well below the last vertebrae) with a sterile razor blade. Blood samples were taken at the following time points: 0min – basal before restraint, 15min and 30min while in the restrainer, and 60min and 120min after being released back into the home cage. Blood was kept on ice through the restraint and then centrifuged to remove plasma. Plasma was kept at −20°C until processed with a I-125 radioimmunoassay (RIA) kit from MP Biomedicals. Duplicates were run for each sample for technical replication.

### Forced swim test (FST)

The modified FST (6, 39) was conducted in regular lighting 1-4 hours after the beginning of the light cycle. Rats were placed in a Plexiglas cylinder measuring 61cm deep with a 19cm diameter filled to 40cm with tap water at 24-26°C. Behavior was analyzed by recording whether the rat was swimming, climbing, diving, or immobile using Kinoscope software (40) by an experimenter blind to genotype.

### Auditory Fear Conditioning

We used the Quirk et al, 2000 (20) method for auditory fear conditioning. Rats were placed into sound attenuated chambers (Med Associates Fairfax VT) with a 33×28×25 cm interior chamber within a 63×45×58 cm sound attenuating outer box and aluminum walls and aluminum rod floor. The fear conditioning paradigm was controlled by Ethovision software (Noldus Information Technology) and consisted of 3 days. On day one, acquisition, rats were allowed to habituate to the chamber for 5 minutes followed by 5 30s tones paired with shock, 0.5mA for.5s, where administered with 3 min inter-trial intervals (ITI). Rats were removed to their homecage and returned to the chamber 24hrs later. For day 2 extinction, after another 5min habituation, 20 tones, 30s with 3min ITIs, were played with no shock at termination. Rats were again returned to the homecage for 24hrs. The last day, day 3 extinction recall, consisted again of a 5min habituation and 20 tones, 30s with 3min ITIs. Freezing, the complete cessation of all movement other than respiration was measured by Freezescan software (CleverSys, Inc) during the 30s tones.

### Statistical analysis

Data are expressed as mean +/- standard error of the mean (SEM). Student’s two-tailed T-test was used to analyze behavioral data and organ weights. A two-way repeated measures analysis of variance (ANOVA) was used to analyze weight, and fear conditioning, and corticosterone data. Fisher’s LSD was used for a priori planned comparisons and post hoc analyses between genotypes. No data points were excluded as outliers. Behavioral data was scored by a researcher blinded to GRKO condition. Sigmaplot (Systat Software) was used to analyze the data. For behavioral experiments, sample size was dependent on the outcome of breeding. Target ‘n’ was 10/group, based on previous power analyses performed in our group; however, ‘n’s were decreased due to missed or ineffective viral injections in the GRKO groups. Effect sizes were calculated to assess the strength of our findings in the face fo reduced ‘n’s’. Effect sizes were in the in the small to medium (pη^2^, ANOVA) and medium to large (Cohen’s d, t-tests) range. 15

## Acknowledgments

J Herman is supported by NIH grants (R01 MH049698, R01 MH101729). R Moloney is supported by a NARSAD Young Investigator Award from the Brain and Behavior Research Foundation. J Scheimann is supported by University of Cincinnati University Research Council Student Faculty Collaboration grant. Y Hu is supported by the Transgenic Core Director Endowment Fund from Cincinnati Children’s Research Foundation. We would also like to thank members of the Herman lab for technical assistance and the Transgenic Animal and Genome Editing Core at Cincinnati Children’s Hospital Medical Center for assistance in generating the floxed *Nr3c1* rat.

## Disclosures

JRS, RDM, PM, RLM, MF, OH, BM, YCH, JPH declare no conflicts of interests.

## Supplementary information

The individual editing activity was validated in rat C6 glioma cell line by T7E1 assay, compared side-by-side with a control sgRNA targeting rat *ApoE*. Cells were transfected with individual sgRNAs (Supplementary Table 1 lists the sgRNAs tested) and cultured for two days. Cells were then harvested for DNA extraction and T7E1 assay. The editing activity was calculated as percentage of the cut band intensity over the total band intensity. The data were represented as relative fold change respective to the control (Supplementary Figure 1).

### Off-target sequences

Off-target sequences associated with each sgRNA that were used to design the SD:nr3c1^fl/fl^ rat were carefully considered (Supplementary Table 2). The required primers (Supplementary Table 3) were designed with Primer3 software Web for the sequences with highest off-target score, based on the CRISPOR Webtool (http://crispor.tefor.net). PCR products were purified with Thermo Scientific™ GeneJET™ Gel Extraction Kit and sequenced by the DNA Sequencing Core at Cincinnati Children’s Hospital Medical Center. We performed off-target analysis on the generated chromatograms by comparing peak-to-peak and nucleotide-to-nucleotide signatures between SD:nr3c1^wt^ and SD:nr3c1^fl/fl^ rats. We observed no off-target events among the most likely off-target sites (Supplementary Table 2).

### Behavioral characterization

GR floxed Heterozygotes from the N2 generation were bred together producing 3 genotypes, SD:nr3c1^fl/fl^ (f/f) relative to SD:nr3c1^fl/-^ (het), and SD:nr3c1^wt^ (wt). Only f/f and wt littermate controls were used for behavioral and physiological measures. All litters were born within 4 days of one another and were 8 weeks of age at the beginning of behavioral assays. Rats were single housed during behavioral experiments in a temperature and humidity controlled vivarium on a 12/12 hour light cycle with ad libitum access to food and water. (n = 12 female f/f and n=12 female wt; n = 12 male f/f and 12 male wt).

Elevated plus maze (EPM) and Open field (OF) tests were performed in dim light 1-4 hours after the beginning of the light cycle. For the EPM, rats were placed on to an elevated plus maze with 2 closed arms ipsilateral from one another and 2 ipsilateral open arms. Arms measured 10cm x 50cm and were elevated 62cm from the ground. Rats were allowed to freely explore the maze for 5 minutes where time spent in open arms and closed arms was analyzed by Ethovision software (Noldus Information Technology, Wageningen - The Netherlands). For the OF, rats were placed in an uncovered plastic square arena measuring 92×92cm and allowed to explore freely for 5 minutes. The number of entries into a square center area measuring 45cmx45cm was analyzed, along with total locomotion, by Ethovision software (Noldus Information Technology, Wageningen - The Netherlands). Animals also underwent the forced swim test (FST) as described in the main text.

The EPM is a common test for anxiety like behavior. Time spent exploring open arms is considered as less anxious, while more anxious animals tend to stay in closed arms where there are less environmental threats. Male and female rats from f/f or wt groups all spend equal amounts of time exploring open arms [Male T(20) = −1.207; p = 0.241(Supplemental Figure 2A); Female T(19) = −0.0939; p = 0.926 (Supplemental Figure 2B)].

Similarly, the OF is a test for anxious behavior with anxious rats spending the majority of the 5min OF test around the periphery where the rodent feels less exposed to environmental dangers. Less anxious rats will explore the center of the arena more. Any alterations to locomotion in animal models can also be seen in the OF test. When tested in the OF, GR floxed rats explored the center for the same amount of time as wt controls in both males [T(20) = 0.762; p = 0.455 (Supplemental Figure 2C] and females [T(19) = −1.563; p= 0.134 (Supplemental Figure 2D)]. The f/f male rats and wt male rats showed no differences in locomotion in the maze [T(20) = 1.365; p=0.187 (Supplemental Figure 2E)], and the same was true for the female f/f and female wt rats [T(19) = −0.0326; p = 0.974 Supplemental figure 2D]. In studies focusing on the effects of stress, the FST is often used as an assay to test for the efficacy of antidepressants to attenuate phenotypes (39). Antidepressants have been shown to decrease immobility, considered passive coping behavior. Further the FST tests for the rodents coping mechanism and how that can change after a manipulation such as chronic stress (2, 4). There were no differences in FST immobility for male [T(20) = 1.960; p = 0.0641 (Supplemental Figure 2G)] nor female f/f and wt rats [T(19) = 1.178; p = 0.253 (Supplemental Figure 2H)].

### Physiology measures

To verify that the gene editing protocol had no effect on periphery physiology, animals were weighed either weekly or bi-weekly from weaning and heart, thymus, and adrenals were dissected from animals after euthanasia (rapid decapitation) and weighed. No differences were observed in body weight profiles in either male (Supplemental Figure 3A) or female (Supplemental Figure 3B) SD:nr3c1^fl/fl^ relative to SD:nr3c1^wt^ controls. Similarly, weight of peripheral organs (thymus, adrenals, heart) were no affected by gene targeting (Supplemental Table 4).

## Supplementary figures

**Supplementary Table 1.**
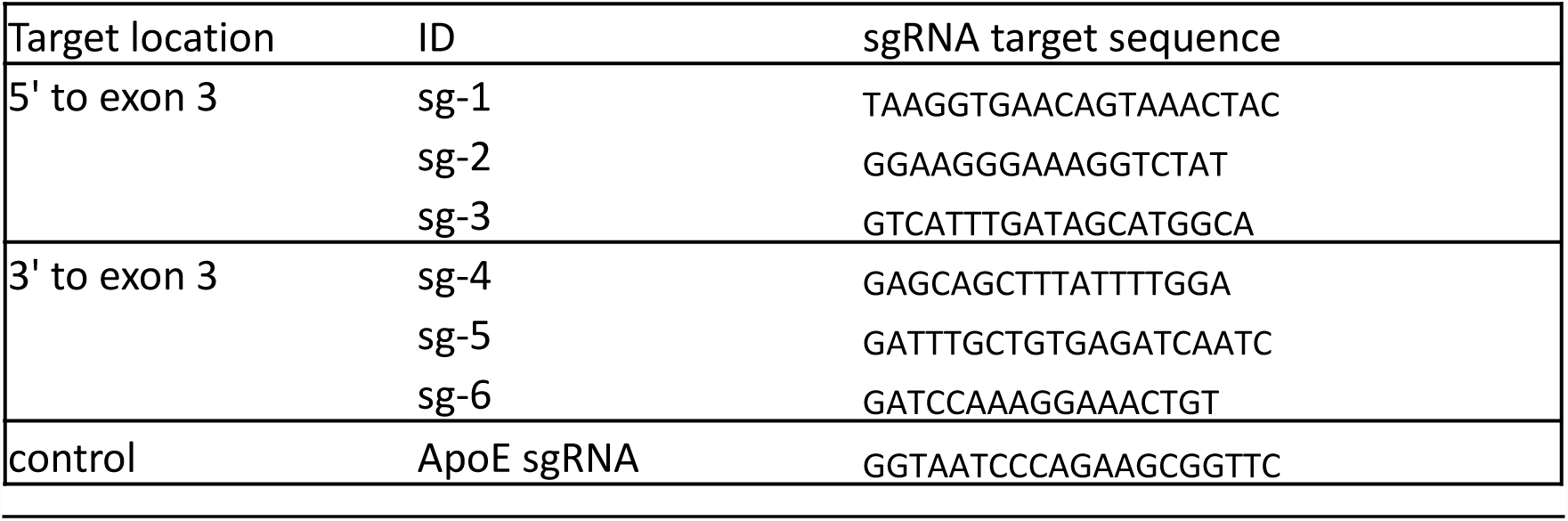
sgRNA target sequence. List of sgRNA chosen to target exon 3 of the *Nr3c1* gene (glucocorticoid receptor) for insertion of loxP sites.

**Supplementary Figure 1.**
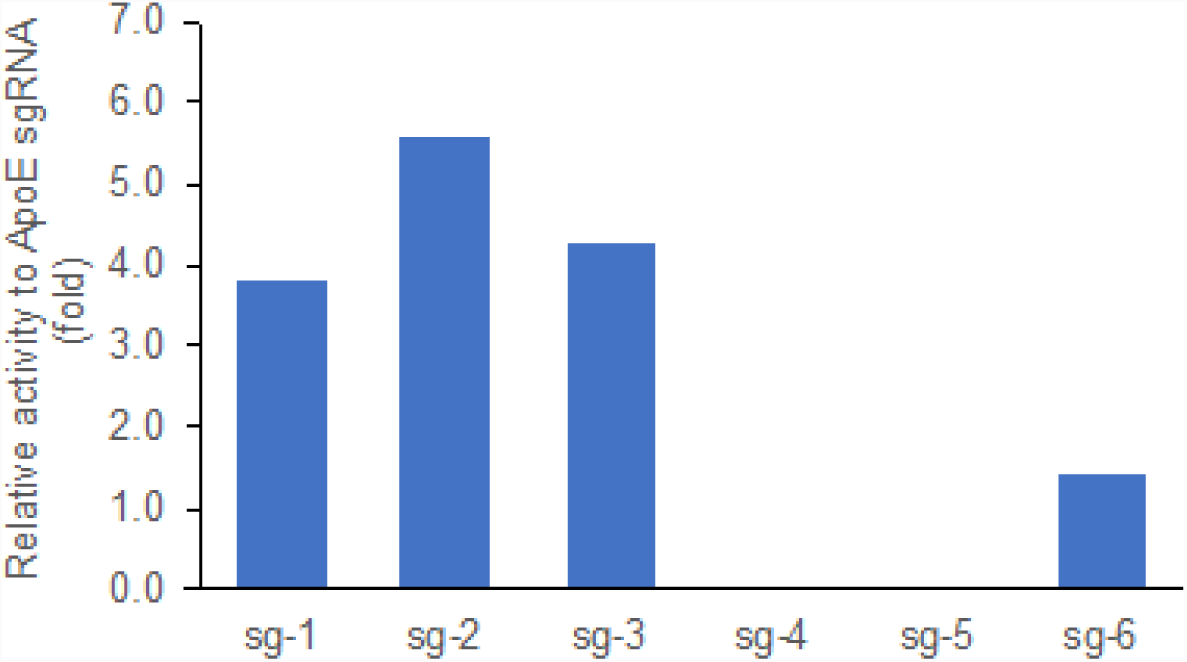
ApoE activity. Validation of sgRNA editing activity by the T7E1 assay in rat C6 glioma cells, compared side-by-side with *ApoE* sgRNA that was previously shown to work efficiency in rat zygotes. Data presented as relative fold change between the percentage of cut band intensity over total band intensity induced by the individual sgRNAs and that of the control ApoE sgRNA.

**Supplementary Table 2.**
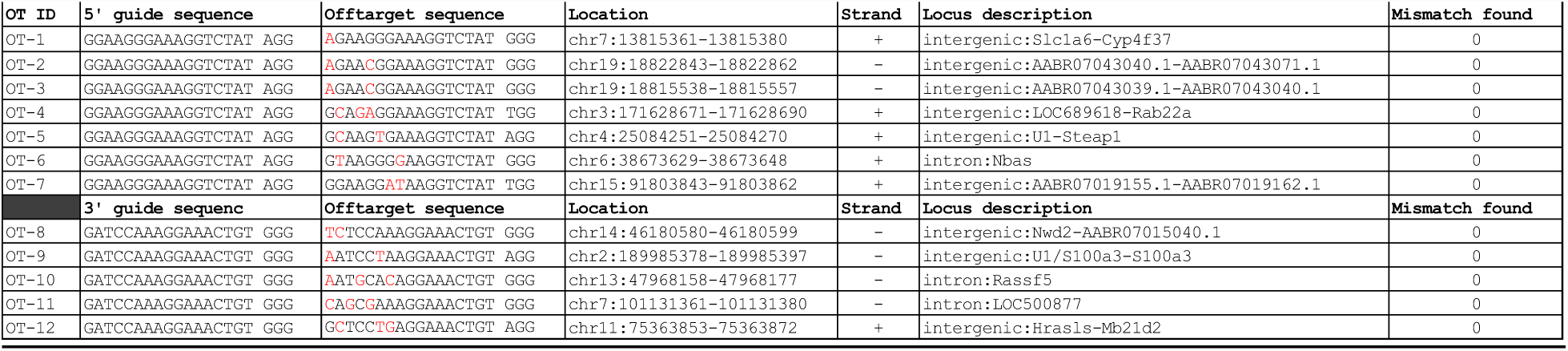
Off-target sequences. List of off-target sequences used to identify the off-target events.

**Supplementary Table 3.**
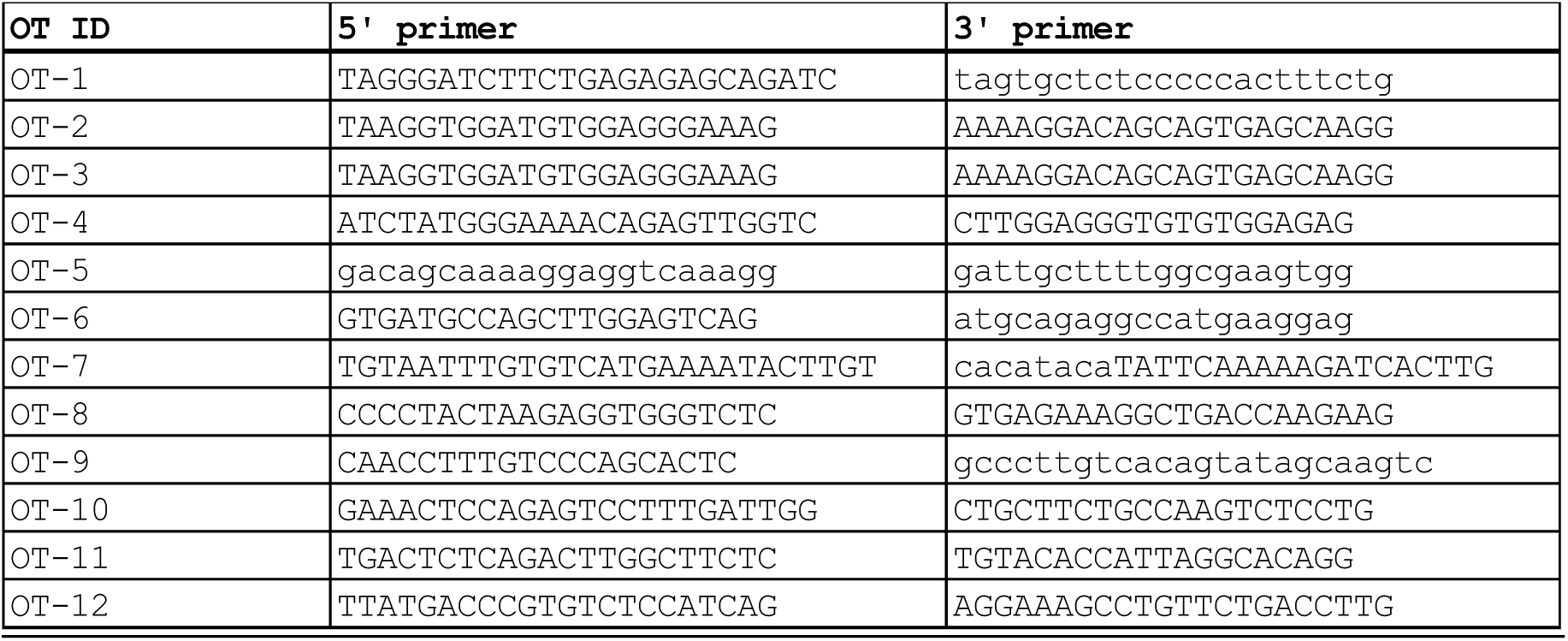
primer sequences. List of primers used to identify the off-target events.

**Supplementary Figure 2.**
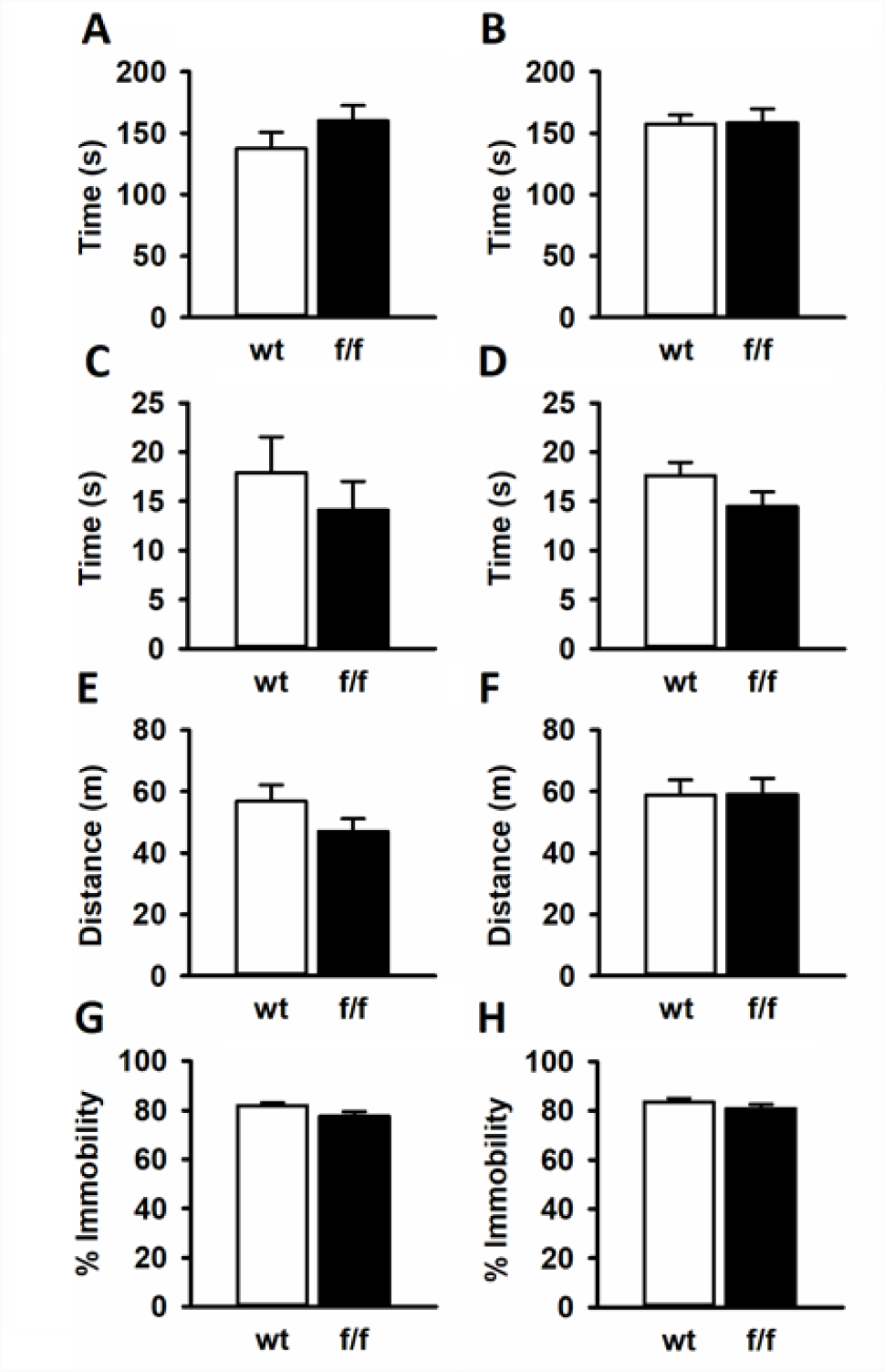
Behavioral characterization. Floxed (f/f) rats did not show behavioral differences from controls in common behavioral assays. Male (A) and female (B) f/f and wt controls spent equal time in the open arms of the elevated plus maze [Males: T(20) = −1.207; p = 0.241; wt mean = 137.573; SEM = 13.24 n = 12; f/f mean = 159.964 SEM 12.599 n = 10; Females: T(19) = −0.0939; p = 0.926; wt mean = 157.213 SEM = 7.502 n = 12; f/f mean = 158.436 SEM = 11.259 n = 9]. Males (C) and females (D) showed no differences in time spent in the center of the open field [Males: T(20) = 0.762; p = 0.455; wt mean = 17.926 SEM = 3.636 n = 9; f/f mean = 14.096 SEM = 2.940 n = 12; Females: T(19) = −1.563; p = 0.134; wt mean = 14.176 SEM = 1.495 n =13; f/f mean = 17.627 SEM = 1.337 n = 9]. No locomotor phenotype was observed in male (E) and female (F) f/f rats compared to wt controls in the open field [Males: T(20) = 1.365; p = 0.187; wt mean = 5688.043; SEM = 528.463 n = 9; f/f mean = 4697.692 n = 12; p = 415.163; Females: T(19) = −0.0326; p = 0.974; wt mean = 5883.908 SEM = 488.249 n = 13; f/f mean = 5907.780 SEM = 516.303 n=9]. There were no differences in immobility in the forced swim test for male (G) or female (H) f/f rats compared to controls [Male: T(20) = 1.960; p = 0.064; wt mean = 81.923 SEM = 1.221 n = 12; f/f mean = 77.583 SEM = 1.938 n = 10; Female: T(19) = 1.178; p = 0.253; wt mean = 83.544 SEM = 1.478 n = 12; f/f mean = 80.854 SEM = 1.751; n = 9].

**Supplementary Figure 3.**
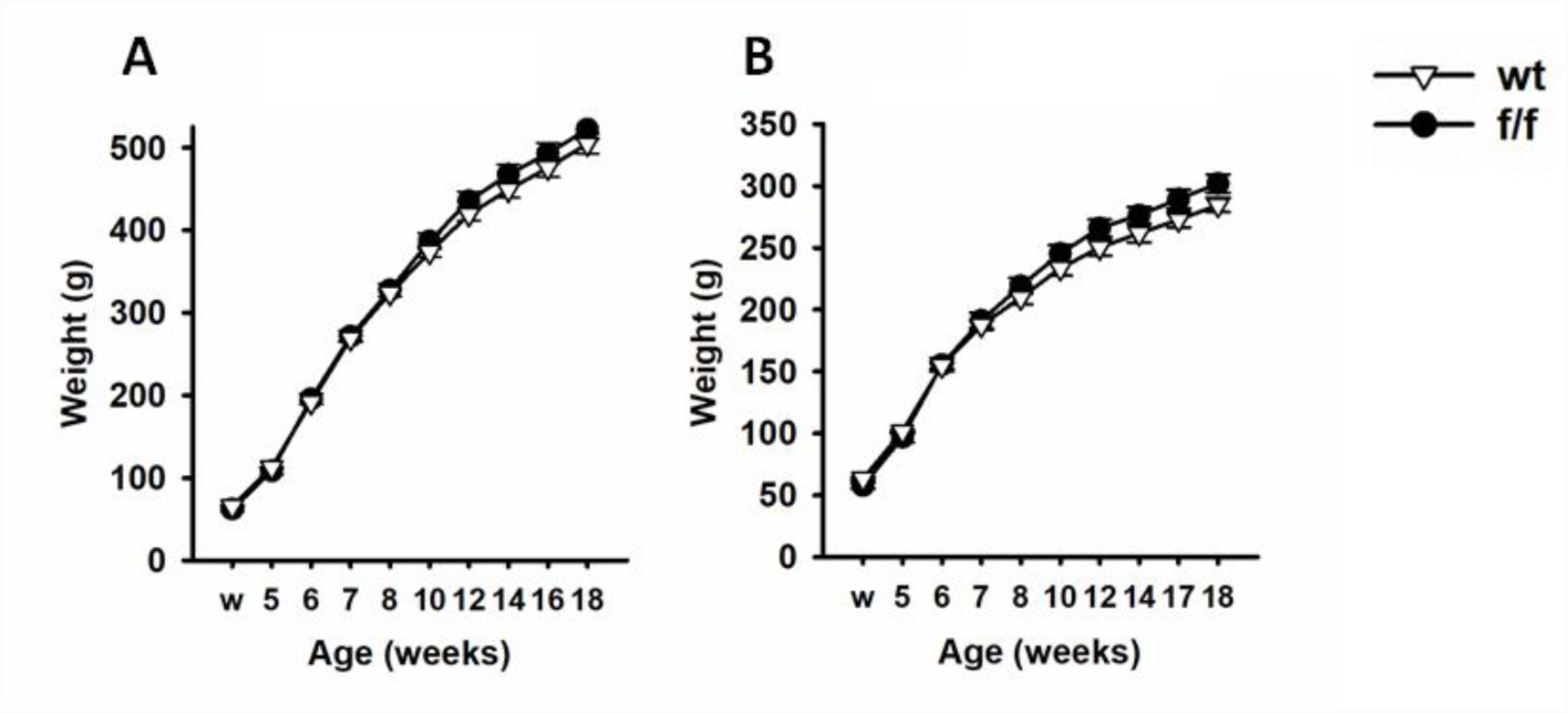
Bodyweight. To ensure floxing GR did not change peripheral physiological measures, rats were weighed weekly. Male (A) and female (B) fl/fl and wt rats did not differ in weight at weening (w) or over the course of the experiment [Male: genotype x time F(1,229) = 0.528; p = 0.475; wt mean = 318.642 SEM = 7.823 n = 13; f/f mean = 327.266 SEM = 8.919 n = 10; Female: genotype x time F(1, 99) = 0.744; p = 0.400; wt mean = 203.203 SEM = 5.226 n = 11; f/f mean = 209.925 SEM = 5.226 n = 9].

**Supplementary Table 4.**
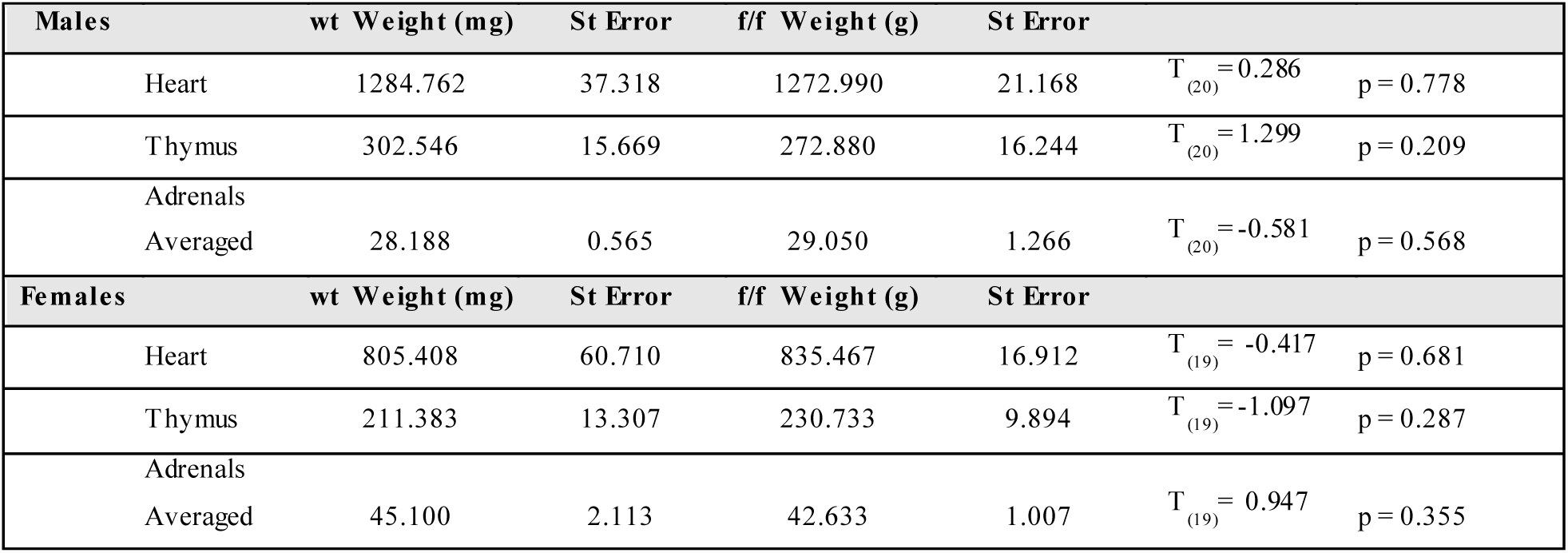
Hearts, thymi, and adrenal weights did not differ in fl/fl vs. wt controls.

## References

1. Herman JP (1993) Regulation of adrenocorticosteroid receptor mRNA expression in the central nervous system. Cell Mol Neurobiol 13(4):349–372.

2. McKlveen JM, et al. (2013) Role of prefrontal cortex glucocorticoid receptors in stress and emotion. Biol Psychiatry 74(9):672–679.

3. McKlveen JM, et al. (2016) Chronic Stress Increases Prefrontal Inhibition: A Mechanism for Stress-Induced Prefrontal Dysfunction. Biol Psychiatry 80(10):754–764.

4. Ghosal S, Bundzikova-Osacka J, Dolgas CM, Myers B, & Herman JP (2014) Glucocorticoid receptors in the nucleus of the solitary tract (NTS) decrease endocrine and behavioral stress responses. Psychoneuroendocrinology 45:142–153.

5. Solomon MB, et al. (2014) The selective glucocorticoid receptor antagonist CORT 108297 decreases neuroendocrine stress responses and immobility in the forced swim test. Horm Behav 65(4):363–371.

6. Wulsin AC, Herman JP, & Solomon MB (2010) Mifepristone decreases depression-like behavior and modulates neuroendocrine and central hypothalamic-pituitary-adrenocortical axis responsiveness to stress. Psychoneuroendocrinology 35(7):1100–1112.

7. Nahar J, et al. (2015) Rapid Nongenomic Glucocorticoid Actions in Male Mouse Hypothalamic Neuroendocrine Cells Are Dependent on the Nuclear Glucocorticoid Receptor. Endocrinology 156(8):2831–2842.

8. Solomon MB, et al. (2015) Neuroendocrine Function After Hypothalamic Depletion of Glucocorticoid Receptors in Male and Female Mice. Endocrinology 156(8):2843–2853.

9. Solomon MB, et al. (2012) Deletion of forebrain glucocorticoid receptors impairs neuroendocrine stress responses and induces depression-like behavior in males but not females. Neuroscience 203:135–143.

10. Whishaw IQ & Tomie J-A (1996) Of Mice and Mazes: Similarities Between Mice and Rats on Dry Land But Not Water Mazes. Physiology & Behavior 60(5):1191–1197.

11. Parker CC, et al. (2014) Rats are the smart choice: Rationale for a renewed focus on rats in behavioral genetics. Neuropharmacology 76 Pt B:250–258.

12. Ellenbroek B & Youn J (2016) Rodent models in neuroscience research: is it a rat race? Dis Model Mech 9(10):1079–1087.

13. Colacicco G, Welzl H, Lipp HP, & Würbel H (2002) Attentional set-shifting in mice: modification of a rat paradigm, and evidence for strain-dependent variation. Behav Brain Res 132(1):95–102.

14. Wang H, La Russa M, & Qi LS (2016) CRISPR/Cas9 in Genome Editing and Beyond. Annu Rev Biochem 85:227–264.

15. Shao Y, et al. (2014) CRISPR/Cas-mediated genome editing in the rat via direct injection of one-cell embryos. Nat Protoc 9(10):2493–2512.

16. Ran FA, et al. (2013) Genome engineering using the CRISPR-Cas9 system. Nat Protoc 8(11):2281–2308.

17. Fu Y, Sander JD, Reyon D, Cascio VM, & Joung JK (2014) Improving CRISPR-Cas nuclease specificity using truncated guide RNAs. Nat Biotechnol 32(3):279–284.

18. Amat J, et al. (2005) Medial prefrontal cortex determines how stressor controllability affects behavior and dorsal raphe nucleus. Nat Neurosci 8(3):365–371.

19. TF G & S M (2015) The Role of the Medial Prefrontal Cortex in the Conditioning and Extinction of Fear. Front Behav Neurosci 9(298).

20. Quirk GJ, Russo GK, Barron JL, & Lebron K (2000) The role of ventromedial prefrontal cortex in the recovery of extinguished fear. J Neurosci 20(16):6225–6231.

21. Quirk GJ, Garcia R, & González-Lima F (2006) Prefrontal mechanisms in extinction of conditioned fear. Biol Psychiatry 60(4):337–343.

22. Ma Y, et al. (2014) Heritable multiplex genetic engineering in rats using CRISPR/Cas9. PLoS One 9(3):e89413.

23. Wood M, et al. (2018) Infralimbic prefrontal cortex structural and functional connectivity with the limbic forebrain: a combined viral genetic and optogenetic analysis. Brain Struct Funct.

24. Boyle MP, Kolber BJ, Vogt SK, Wozniak DF, & Muglia LJ (2006) Forebrain glucocorticoid receptors modulate anxiety-associated locomotor activation and adrenal responsiveness. J Neurosci 26(7):1971–1978.

25. Hartmann J, et al. (2017) Forebrain glutamatergic, but not GABAergic, neurons mediate anxiogenic effects of the glucocorticoid receptor. Molecular Psychiatry 22:466–475.

26. Zhang XH, Tee LY, Wang XG, Huang QS, & Yang SH (2015) Off-target Effects in CRISPR/Cas9-mediated Genome Engineering. Mol Ther Nucleic Acids 4:e264.

27. Fu Y, et al. (2013) High-frequency off-target mutagenesis induced by CRISPR-Cas nucleases in human cells. Nat Biotechnol 31(9):822–826.

28. Cho SW, et al. (2014) Analysis of off-target effects of CRISPR/Cas-derived RNA-guided endonucleases and nickases. Genome Res 24(1):132–141.

29. Morgan MA & LeDoux JE (1995) Differential contribution of dorsal and ventral medial prefrontal cortex to the acquisition and extinction of conditioned fear in rats. Behav Neurosci 109(4):681–688.

30. Marquis J, Killcross S, & Haddon J (2007) Inactivation of the prelimbic, but not infralimbic, prefrontal cortex impairs the contextual control of response conflict in rats. European Journal of Neuroscience 25:559–566.

31. Bielajew C, et al. (2003) Strain and gender specific effects in the forced swim test: effects of previous stress exposure. Stress 6(4):269–280.

32. Fujisaki C, et al. (2003) An immnosuppressive drug, cyclosporine-A acts like anti-depressant for rats under unpredictable chronic stress. J Med Dent Sci 50(1):93–100.

33. Molina VA, Heyser CJ, & Spear LP (1994) Chronic variable stress or chronic morphine facilitates immobility in a forced swim test: reversal by naloxone. Psychopharmacology (Berl) 114(3):433–440.

34. Breslau N, Davis GC, Andreski P, Peterson EL, & Schultz LR (1997) Sex differences in posttraumatic stress disorder. Arch Gen Psychiatry 54(11):1044–1048.

35. Breslau N & Anthony JC (2007) Gender differences in the sensitivity to posttraumatic stress disorder: An epidemiological study of urban young adults. J Abnorm Psychol 116(3):607–611.

36. Kessler RC, Sonnega A, Bromet E, Hughes M, & Nelson CB (1995) Posttraumatic stress disorder in the National Comorbidity Survey. Arch Gen Psychiatry 52(12):1048–1060.

37. McLean CP, Asnaani A, Litz BT, & Hofmann SG (2011) Gender differences in anxiety disorders: prevalence, course of illness, comorbidity and burden of illness. J Psychiatr Res 45(8):1027–1035.

38. Madisen L, et al. (2015) Transgenic mice for intersectional targeting of neural sensors and effectors with high specificity and performance. Neuron 85(5):942–958.

39. Cryan JF, Valentino RJ, & Lucki I (2005) Assessing substrates underlying the behavioral effects of antidepressants using the modified rat forced swimming test. Neurosci Biobehav Rev 29(4-5):547–569.

40. Kokras N, Baltas D, Theocharis F, & Dalla C (2017) Kinoscope: An Open-Source Computer Program for Behavioral Pharmacologists. Front Behav Neurosci 11:88.

